# Texture Analysis of Dystrophic Muscle Biopsies

**DOI:** 10.1101/273771

**Authors:** Wlodzimierz Klonowski, Bozenna Kuraszkiewicz, Anna M. Kaminska, Anna Kostera-Pruszczyk

## Abstract

Despite the introduction of a full range of genetic diagnostic tests and sophisticated techniques in modern pathology, interpretation of histopathological images obtained from muscle biopsies remains important in the daily practice of neuropathology since it can give indications of the severity and the rate of progression of neuromuscular disease. In this paper, we propose a simple and time saving method for quantitative assessment of severity of Duchenne Muscular Dystrophy (DMD) based on computer-aided analysis of histopathological images obtained from biopsies of dystrophic muscles. Using this method, **c**olour **f**iltration **p**ixel-by-**p**ixel of the whole virtual slides (*CFPP method*) is adopted to semi-quantitative evaluation of morphological structure of the muscular tissue. Results demonstrate usefulness of the proposed method in neuropathological assessement of DMD severity.

## 1 Introduction

Recent developments of automated systems for analysis of genetic material have caused that nowadays histopathological methods based on images from muscle biopsies are applied not so often as in previous decades, since they are more time-consuming. Nevertheless, histopathological evaluation of muscle biopsies are still clinically applied along with other diagnostic methods as muscle MRI or 6 minutes walk test. The aim of this study is to develop simple and quick semi-automatic method for analysis of histopathological images to assist evaluation of muscle dystrophies, especially Duchenne Muscular Dystrophy (DMD). The method is based on computer-aided analysis of images - **c**olour **f**iltration **p**ixel-by-**p**ixel (*CFPP method*) of the whole histopathological virtual slides.

DMD is a severe genetic disease characterized by a progressive muscle weakness and loss in structural integrity of muscle fiber membranes, caused by deficiency of the dystrophin protein [1]. Respiratory complications and cardiomyopathies are common causes of DMD patients premature death.

Healthy muscle has a characteristic appearance, and is made up of closely-packed fibers, which are more or less evenly sized. Dystrophic changes on muscle biopsy include variation in fiber size, enlarged internal nuclei, grown fatty and connective tissue [2], and presence of regenerating and degenerating fibers [3]. As the disease progresses the microscopic images will show a greater proportion of fatty and connective tissue. In later pathology the image shows an increase in internal nuclei and variability of myofiber size [4]. Hence, digital pathology helps in neuropathological diagnostics [5].

## 2 Materials and Methods

Among many features, severity of dystrophic muscles is assessed based on the amount and distribution of irregular myocytes, and the amount of fatty and connective tissue in the microscopic images obtained from biopsy (*whole specimen*). When grading the severity of dystrophy, pathologist concentrates his/her attention on some fragments of the image. The evaluation based on the whole specimen is frequently compared to the evaluation in the manually chosen and magnified *regions of interests (ROIs)* containing *objects* (cells’ nuclei) that are hematoxilin-stained (H) and so turned blue, and *objects* (myocytes) that are eosin-stained (E) and so turned pink.

Manual grading, based on visual assessment, is frequently time consuming. Besides, it is usually focused on up to 100-200 myofibers being assessed along with surrounding connective tissue. This kind of visual evaluation requires patience and accuracy in choosing *ROIs*. Hence, our objective is to develop method for quick and global assessment of microscopic images of dystrophic muscles.

Among the full range of methods used in the muscle dystrophy analysis earlier we choose Higuchi’s fractal dimension algorithm (cf. [6]). However, in analysis of biopsies of DMD muscles the results of such fractal analysis are not satisfactory - the method does not differentiate between moderate and advanced stages of the illness (Fig.1.).

**Fig. 1.**
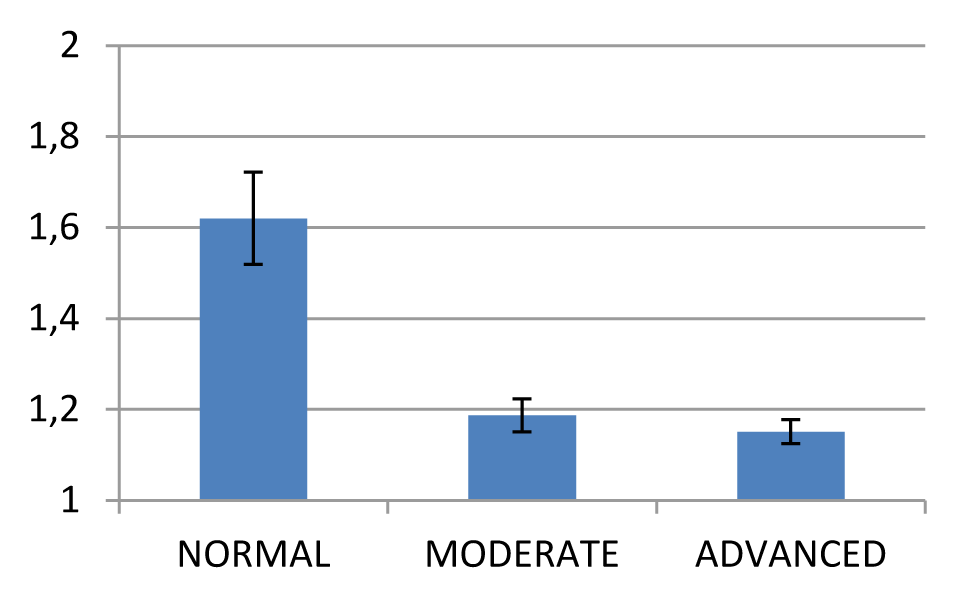
Mean±SD values of Higuchi fractal dimension calculated from microscopic images of biopsies of normal muscle tissue (n=3) and of moderate (n=8) and advanced DMD (n=5).

Herein, we propose a new dedicated method for quick semi-automatic assessment of severity (stage) of DMD from dyes-stained microscopic images of muscle biopsies.

We retrospectively analyzed 13 digital microscopic images of DMD muscle tissue from 4 patients (average±SD age 6.5±1.3 years). The images were obtained from 2 subjects in moderate (8 slides) and 2 subjects in advanced DMD stage (5 slides) at the Department of Neurology, Medical University of Warsaw. The images were acquired with x400 magnification, from original microscopy slides coming from biopsies. The biopsies were performed as a part of medical diagnostic procedure, so no IRB approval was required. Additionally, we have analyzed 3 microscopic slides of normal muscle tissue.

The images were analyzed on a standard PC with Intel i7 dual core processor and 8GB RAM. All analyses were performed in MATLAB 2016b.

Our new method is based on **c**olor **f**iltration **p**ixel-by-**p**ixel of the whole virtual slides (*CFPP method*) and semi-quantitative evaluation of morphological structure of the muscular tissue. The method is simple and quick – it does not require either time consuming manual marking of artifact-free representative *ROIs* or complicated detection of myocytes to grade the dystrophic tissue. Instead, we propose to count the numbers of pixels belonging to the H-stained nuclei (*L*_*H*_), to the E-stained myocytes (*L*_*E*_), and to the E-stained connective tissue including sarcolemma and not stained regions belonging to adipocytes (*L*_*C*_) in the given whole specimen.

We define two quantities characterizing dystrophic changes that will be called the *connective tissue indices*:

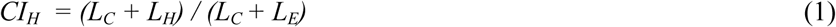

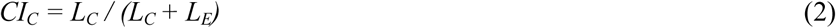

The calculations are easily done with MATLAB. Each whole specimen is appropriately color filtered pixel by pixel. Color-coding of computer images most often uses RGB system - the color of a pixel is expressed as a triplet, *(r,g,b)* i.e. *(read, green, blue)*, each component of which can vary from *0* for the darkest one to *255* for the brightest one (Fig. 2.).

**Fig. 2.**
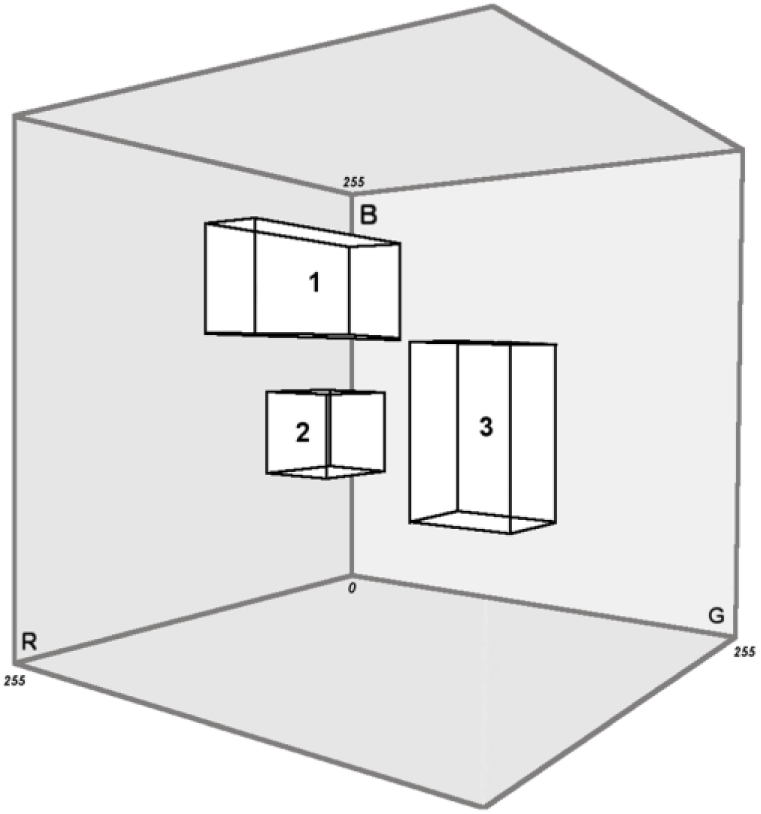
RGB-space and its subspaces corresponding to different colours of pixels.

The whole RGB-space has nearly 17 millions of points (256×256×256). To characterize biopsies of DMD muscles we consider cardinality of two subspaces – one classified as consisting of pixels corresponding to H-stained nclei and another classified as consisting of pixels corresponding to E-stained nuclei.

So, for each of .tiff image to count pixels belonging to the nuclei that are H-stained we choose pixels with *(r,g,b)* components greater than minimum values further denoted with index *Hn,* but smaller than maximum values further denoted with index *HN, i.e.* pixels fulfilling the condition

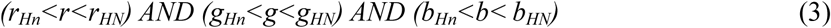

To count pixels belonging to the E-stained connective tissue we choose pixels with *(r,g,b)* components greater than minimum values further denoted with index C*m* but smaller than maximum values further denoted with index *CM, i.e.* pixels fulfilling the condition

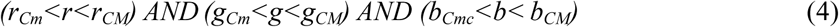

Pixels belonging to adipocytes are ommitted or partially included in the count of connective tissue pixels, as fatty tissue is usually completely washed out from the microscopy specimens.

In general, to count pixels belonging to the E-stained myocytes we choose pixels with *(r,g,b)* components greater than minimum further denoted with index *Em* but smaller than maximum values further denoted with index *EM, i.e.* pixels fulfilling the condition

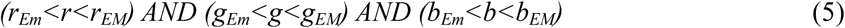

But quite often, like in the presented case, for the number of pixels belonging to E-stained myocytes, *L*_*E*_, one can take all the remaining pixels in the image

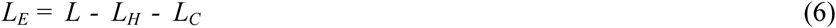

where *L* denotes the total number of pixels in the given specimen.

It is important that for at least one component, *r* or *g* or *b,* the intervals of values for pixels belonging to H-stained (2) and to E-stained (3, 4) structures should be completely disjoint. Such color filtration also filters out the remainings of the empty background and majority of artefacts.

When a fragment of a slide is viewed in MATLAB then clicking on a pixel shows *(r,g,b)* components of this pixel. To choose *Hn* values one clicks on several dark-blue pixels belonging to the nuclei, one writes down the shown *(r,g,b)* values, and for calculation of *L*_*H*_ one takes average value of the corresponding component as the limits in (2); similarly, to choose *HN* values one clicks on several light-blue pixels. To choose *Em* or *Cm* values one clicks on dark pixels belonging to myocytes or connective tissue, respectively; to choose *EM* or *CM* values one clicks on some light pixels belonging to myocytes or connective tissue, respectively. One writes down the shown *(r,g,b)* values and for calculation of *L*_*E*_ or *L*_*C*_ one takes average values of the corresponding component as the limits in (4) and (5) (cf. Fig. 3.).

**Fig. 3.**
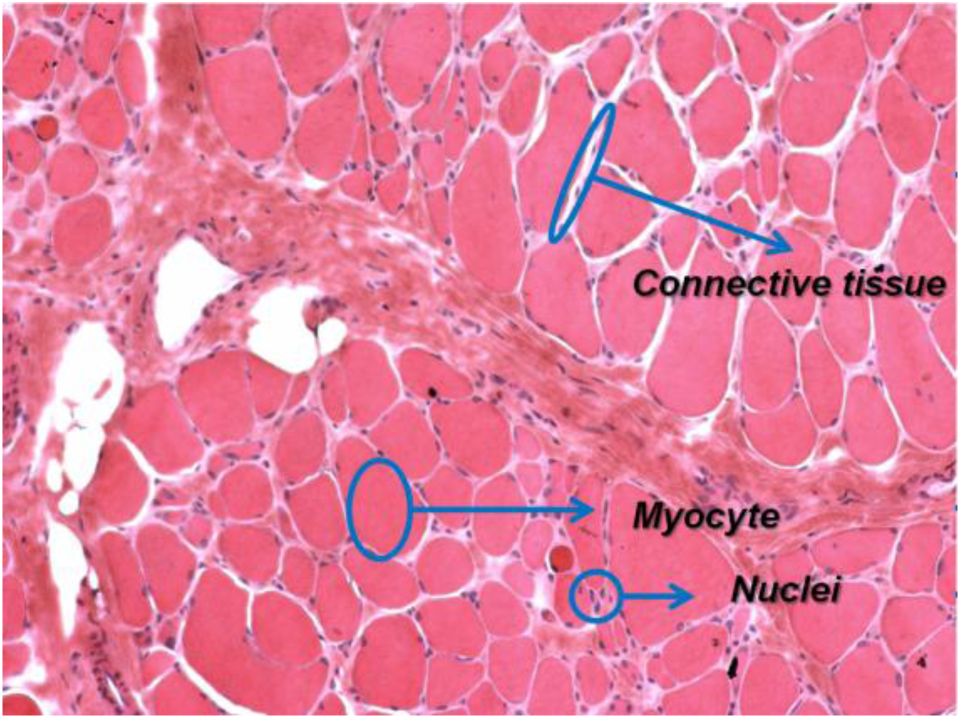
Example of regions: Nuclei - H-stained, counted into *L*_*H*_; Myocyte – E-stained, counted into *L*_*M*_; Connective tissue – partially E-stained sarcolemma with not stained adipocytes, counted into *L*_*C*_. These regions were used to establish the limits of *(r,g,b)* in Ineqs. (7)-(8).

In the presented case, to count the number of pixels belonging to H-stained nuclei, *L*_*H*_, we choose the following *(r,g,b)* intervals:

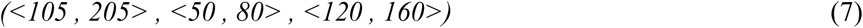

To count the number of pixels belonging to E-stained connective tissue, *L*_*C*_, we choose the following *(r,g,b)* intervals:

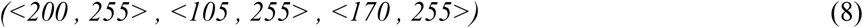

To calculate *L*_*H*_, *L*_*E*_ and *L*_*C*_ the whole specimen may be subdivided into any number of disjoint parts or even read into RAM pixel by pixel. It is important since it makes possible to analyze specimens even on a PC with 8 GB of RAM. *L*_*H*_, *L*_*E*_ and *L*_*C*_ are used to calculate the connective tissue indices, *CI*_*H*_ and *CI*_*C*_ (Eqs. (1)-(2)).

## 3 Results

We have calculated connective tissue indices *CI*_*H*_ and *CI*_*C*_ for 12 slides of 4 patients in two grades of DMD and for 3 images of normal muscles. Examples are shown on Figs. 4.-6. and mean values and standard deviation of the connective tissue indices moderate and advanced DMD compared to normal muscle are shown on Figs.7.-8.

**Fig. 4.**
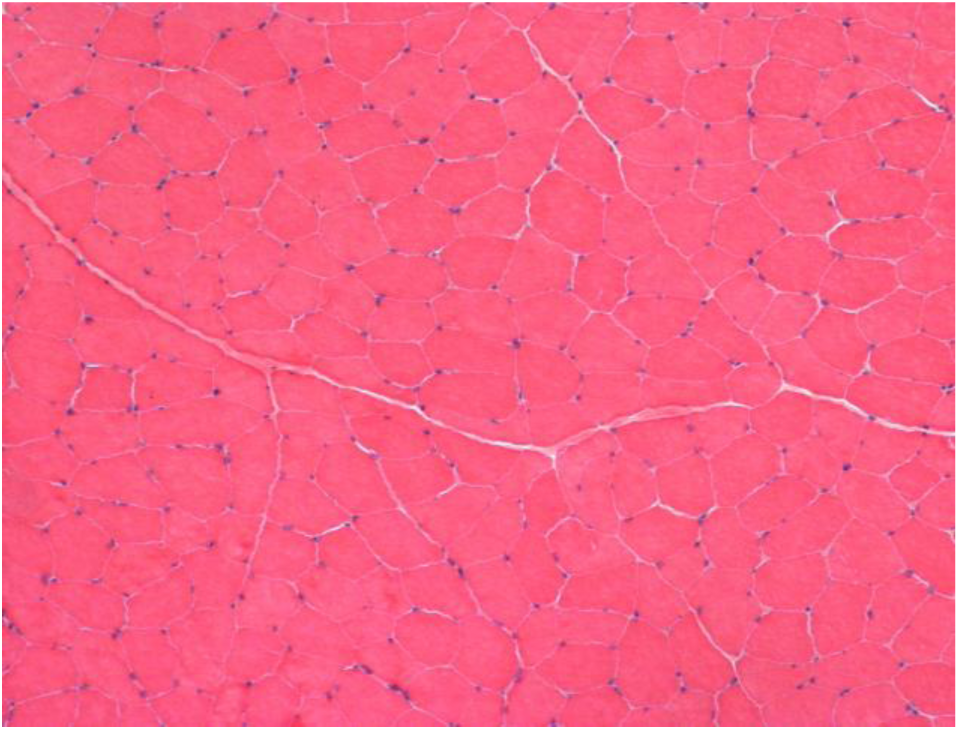
Normal muscle. *CI_H_ =* 0.032 ± 0.001.

**Fig. 5.**
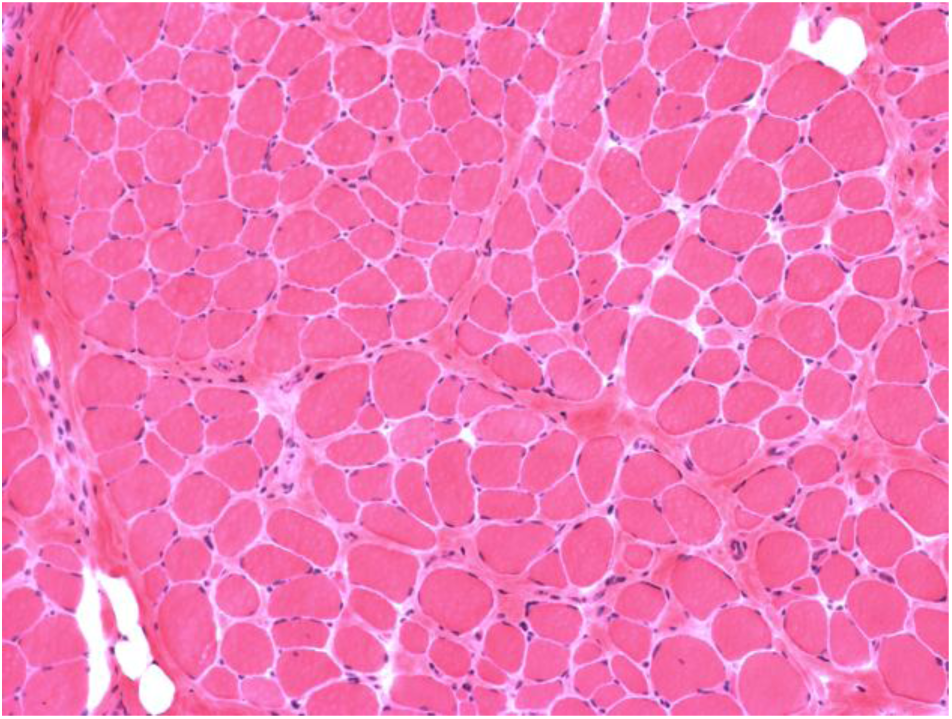
Moderate stage of DMD, *CI_H_* = 0.18 ± 0.03.

**Fig. 6.**
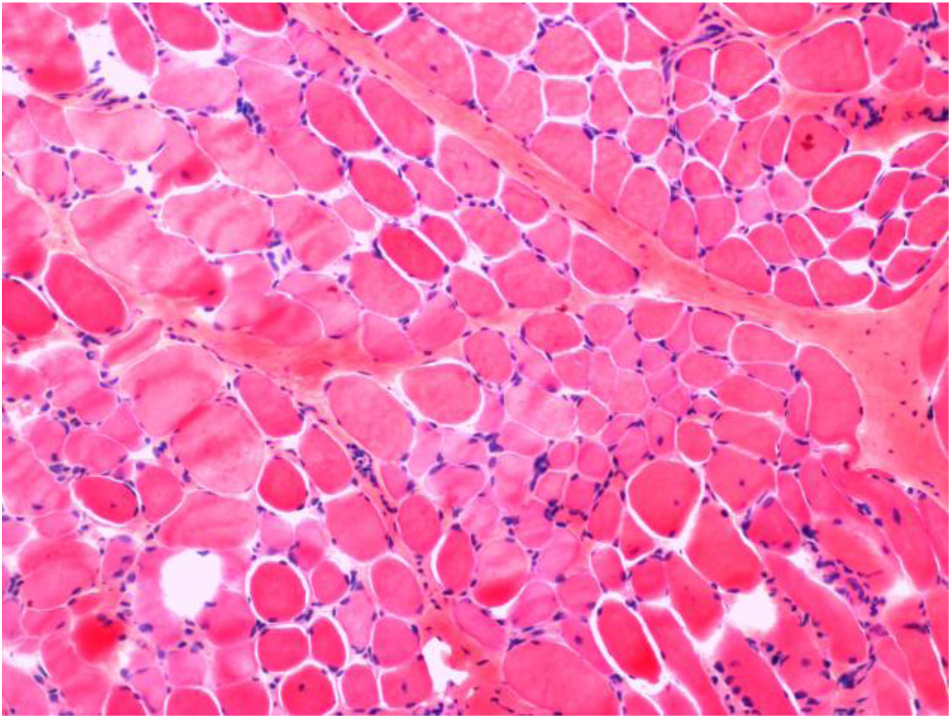
Advanced stage of DMD, *CI_H_ =* 0.31 ± 0.09.

**Fig. 7.**
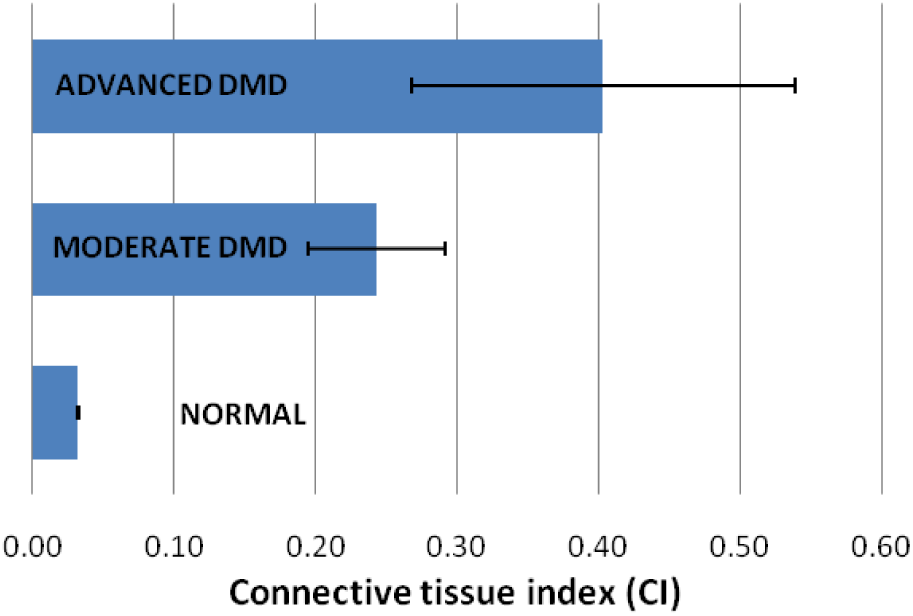
Connective tissue indices *CI*_*H*_ for the analyzed DMD and normal muscle tissue images.

**Fig. 8.**
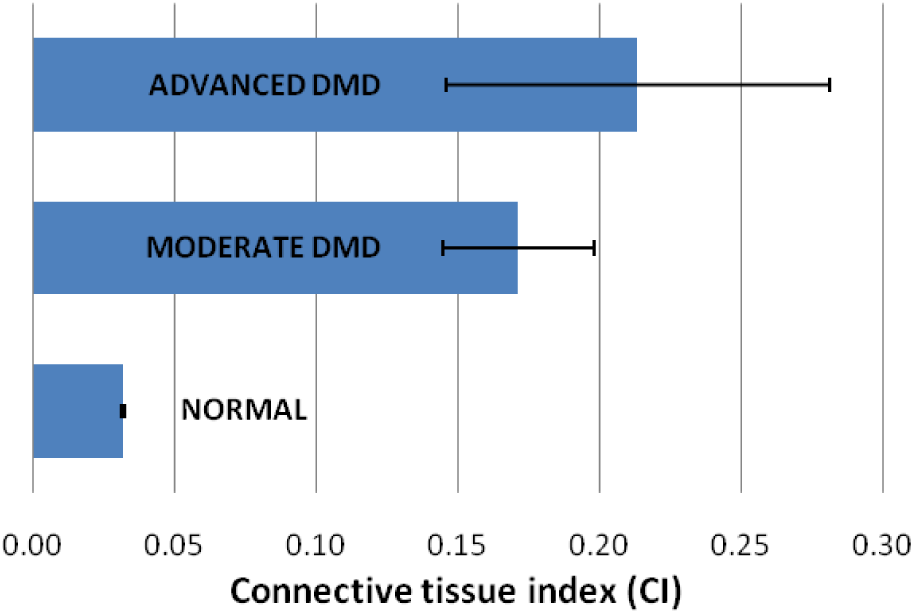
Connective tissue indices *CI*_*C*_ for the analyzed DMD and normal muscle tissue images.

## 4 Conclusions

We have presented a fast semi-automatic method for evaluation of muscle tissue microscopy images. CFPP method is simple and allows to distinguish DMD tissue from normal muscle tissue, as well as potentially gives a possibility to evaluate the severity of DMD. From the operator’s point, the method requires only choosing of a small representative regions of muscle tissue, fatty and connective tissue, and of nuclei. As the method is applicable globally to a single image, it does not require colour scale standardization or brightness and contrast among sets of microscopic images. Here the method is dedicated to DMD images, but its general principle is also applicable to other histological images (cf. [7]).

Connective tissue indices calculated shown that *CI*_*H*_ index (1) better differentiates stages of DMD than *CI*_*C*_ index (2) (cf. Figs. 7.-8.) but further research is needed. It is worth to highlight that *CI*_*H*_ includes the count of pixels belonging to nuclei regardless whether the nuclei belong to muscle or connective tissue. We also consider the situation, that for some slides *CI*_*C*_ might be a better measure of DMD severity, as those slides might lack of nuclei due to anatomical or pathological conditions.

Presented results are promising and further research is planned on a large data set, including microscopy images from different types of dystrophy. Moreover, initial step assessing the quality of analyzed microscopy images is foreseen to be implemented for CFPP method.

## Acknowledgements

This work was partially supported by Nalecz Institute of Biocybernetics and Biomedical Engineering Polish Academy of Sciences through its statutory activity. We also thank COST Action BM1304 MYO-MRI for interesting discussions.

## References

[1] A.Y. Manzur and F. Muntoni. Diagnosis and new treatments in muscular dystrophies. J Neurol Neurosurg Psychiatry, vol.80, pp.706–714, 2009.

[2] F.M. Norwood et al. EFNS guideline on diagnosis and management of limb girdle muscular dystrophies. Eur J Neurol, vol.14, no.12, pp.1305–1312, 2007.

[3] E.M. McNally and P. Pytel. Muscle diseases: the muscular dystrophies. Annu Rev Pathol, vol.2, pp. 87–109, 2007.

[4] T.A. Wren et al. Three-point technique of fat quantification of muscle tissue as a marker of disease progression in Duchenne muscular dystrophy: preliminary study. AJR Am J Roentgenol, vol.190, pp.W8–W12, 2008.

[5] W. Klonowski, Applications of Chaos Theory Methods in Clinical Digital Pathology, In: Handbook of Applications of Chaos Theory, Ch.H. Skiadas & C. Skiadas, Eds. CRC Press, Boca Raton, New York, 2016, pp.681–690.

[6] R.A. Lerski. J.D. de Certaines, D. Duda, W. Klonowski, G. Yang, J.L. Coatrieux, N. Azzabou, P.-A. Eliat. Application of texture analysis to muscle MRI: 2 – technical recommendations. EPJ Nonlinear Biomedical Physics 3:2, 2015, https://epjnonlinearbiomedphys.springeropen.com/articles/10.1140/epjnbp/s40366-015-0018-0

[7] W. Klonowski, A. Korzynska, R. Gomolka, Computer analysis of histopathological images for tumor grading. Physiological Measurement, prepublished on-line http://iopscience.iop.org/article/10.1088/1361-6579/aaa82c/pdf

